# Cryo-EM Reveals VP1/VP2 N-Terminal Conformations in Adeno-Associated Virus 2 Capsids

**DOI:** 10.64898/2026.10.16.759763

**Authors:** Innocent O. Osuya, Shirin S. Kamalaldinezabadi, Ananya Sengupta, Gwimoon Seo, Michael S. Chapman, Scott M. Stagg

## Abstract

The AAV2 capsid assembles from three viral proteins, VP1, VP2, and VP3, expressed from a single cap gene. In the assembled T=1 60-subunit icosahedral capsid, they have a canonical stoichiometry of 1:1:10 . Relative to VP3, VP1 and VP2 have N-terminal extensions that are required for endosomal escape and nuclear trafficking; however, these regions remain poorly resolved due to low stoichiometry, symmetry mismatches, and conformational heterogeneity of their N-termini. Here, we generated empty AAV2 capsids using a baculovirus expression system with time-course harvests, yielding approximately 2:3:5 and 1:2:7 VP1:VP2:VP3 ratios on days 3 and 5 post-infection, respectively. Cryo-EM single-particle and subparticle reconstructions revealed distinct N-terminal configurations associated with these different VP stoichiometries. Day 3 capsids contained an internal density protruding from the twofold axis and lacked ordered βA preceding residue 237, whereas day 5 capsids lacked the twofold protrusion and instead showed density associated with the fivefold axis, with βA ordered to approximately residues 219 - 221. Fivefold subparticle reconstruction of day 5 capsids further resolved a density plug occluding the pore and an underlying density ring. These findings link VP stoichiometry to distinct conformations of the VP1/VP2 N-termini and provide structural evidence for alternative organization of these regions within the AAV2 capsid, offering new insights into the conformational landscape of AAV capsid heterogeneity.

**Significance:** Endosomal escape and nuclear trafficking are fundamental to adeno-associated virus (AAV) transduction, yet the structural conformations of the N-terminal regions of the capsid proteins (VP1 and VP2), which are critical for these processes, remain poorly understood because of their low stoichiometric abundance, conformational flexibility, and the limitations of icosahedral symmetry averaging. Using a baculovirus expression system to increase the relative stoichiometric abundance of VP1 and VP2, we applied cryo-electron microscopy with local signal subtraction and focused classification to reveal previously uncharacterized structural features, including distinct βA-strand conformations, an internal density protrusion at the twofold symmetry axis, and differences in the density plug at the fivefold pore. These findings advance our understanding of AAV2 capsid structural heterogeneity and provide a foundation for investigating how VP1/VP2 conformational variability relates to capsid function during infection.

## Introduction

Adeno-associated Viruses (AAVs) are nonpathogenic members of the *Parvoviridae* family that require helper functions from other viruses, like adenovirus or herpesvirus, for efficient replication ^1^. The nonpathogenic nature of its infection profile, combined with its broad tropism and sustained gene expression, has made recombinant adeno-associated virus (rAAV) vectors attractive for *in vivo* gene therapy, with clinical success further establishing their therapeutic potential and leading to the regulatory approval of multiple AAV-based therapies by the FDA and EMA for the treatment of diverse genetic and inherited disorders ^2–8^. In addition, more than 225 clinical trials investigating AAV-based therapies for over 30 genetic diseases are currently ongoing ^9^. Despite these advances, many critical questions about the fundamental biology and structural characteristics of AAV trafficking remain unanswered, including the structural mechanisms of endosomal escape and nuclear trafficking, which are associated with the N-terminal sequences of capsid proteins VP1 and VP2.

The AAV genome is a single-stranded DNA molecule approximately 4.7 kb in length, with two open reading frames: the cap and rep ^3,10,11^ genes. The cap gene encodes three viral proteins, VP1, VP2, and VP3, expressed through alternative splicing and start codons. These variants are incorporated into the capsids at a canonical ratio of 1:1:10 and share most of their amino acid sequence, differing only at their N-termini. Specifically, VP2 is approximately 137 amino acids shorter than VP1, while VP3 is approximately 65 residues shorter than VP2 and 202 residues shorter than VP1^12–14^. The N-terminal region of VP1 that is unique to that protein is called VP1u. Although they differ in their N-terminal extensions, VP1, VP2, and VP3 share a common sequence that encodes the body of each protein, and together they assemble to form the AAV capsid, approximately 25 nm in diameter, composed of 60 subunits with T=1 icosahedral symmetry ^3,10,11^.

The structural topology of AAV capsids (Fig. S1) is highly conserved across serotypes, having a depression at the twofold axis, three protrusions surrounding the threefold axes, and 12 narrow pores at the fivefold axes ^15^. Each subunit adopts an nine-stranded β-barrel motif ^16^ containing two antiparallel β-sheets, with interstrand loop insertions comprising approximately 60% of the structure ^15^. These loops contain small regions of β-structure and a conserved α-helix that form the wall of the twofold depressions, while interstrand loops from neighboring subunits near the threefold axes intertwine to form surface spikes. Two short stretches of strands within each subunit form a β-ribbon that clusters at the icosahedral fivefold axis, creating the cylindrical pores that connect the capsid interior to the exterior ^15^. These pores extend outward from the capsid surface and are encircled by a depression lined with a loop between the βH and βI strands of the β-barrel ^15,17^.

Biochemical and mutational evidence shows that the N-terminus of VP1 and VP2 in some parvoviruses can be extruded through the pores at the fivefold axes while the capsid remains intact ^18,19^. The crystal structure of canine parvovirus (CPV) virions supports the hypothesis that the weak density within the fivefold pore corresponds to the N-terminal peptide sequence ^17^ of VP1. In addition, the fivefold pore of the CPV capsid has been postulated to be the site of genome import into the preformed capsid, genome export during uncoating, and exposure of the VP1/VP2 N-terminus during intracellular trafficking ^18,19^. These VP1/2 N-terminal regions are, at best, poorly resolved in icosahedral single-particle reconstructions, presumably due to conformational flexibility and low occupancy ^20^. Structural uncertainties near the twofold and threefold symmetry axes have also been observed, including repositioned β-strands and flexible connecting loops ^21^. The disordered and variable nature of these regions makes detailed structural interpretation difficult, consistent with diverse and perhaps dynamic conformational states within the assembled capsid.

The extrusion of the VP1/VP2 N-termini is critical to the virus life cycle. AAV2 infection begins with the attachment of the capsid to heparan sulfate proteoglycans on the cell surface, followed by docking to the primary receptor, AAVR ^22^. The virion enters by clathrin-dependent or independent endocytosis and is trafficked within the endosomal compartment ^23^. Studies have shown that AAV requires endosomal acidification to escape from the late endosome and traffic to the nucleus ^14,19,23,24^. Before endosomal escape, the virion undergoes conformational changes that expose the N-termini of VP1 and VP2, both of which are required for endosomal escape and nuclear entry, with basic amino acid clusters within VP1u contributing to nuclear import in AAV2 and AAV6 ^14,25,26^. Exposure of the VP1 N-terminus is functionally important because it makes the VP1u phospholipase A2 domain accessible, consistent with the lack of phospholipase activity in intact capsids. Furthermore, heat treatment of virions can induce VP1 N-terminal exposure, providing experimental evidence for this conformational transition ^18^.

The N-termini of VP1 and VP2, proposed to be localized on the inside of assembled capsids, have not been visualized in any parvovirus crystal structure; however, cryo-EM reconstruction studies of AAV2 resolved "globules" inside the twofold axis that were ascribed to these regions ^11^, but, notably, they have not been seen in all studies ^10^. Consistent with a variable organization of the N-termini, cryo-electron tomography of the AAV2-AAVR complex revealed diffuse density near the twofold axis and variable ordering of the βA strand, which were interpreted as evidence for at least two alternate N-terminal conformations ^27^. Variability in the exact presentation of these structural features raised the question of whether uncharacterized sample heterogeneity caused the structural variation, particularly given the diversity of heterologous and native AAV sources used for structural studies ^28–30^.

In the present study, we hypothesize that variable VP1, VP2, and VP3 expression *in vitro* is a leading cause of variability in the appearance of features ascribed to the VP1/2-unique regions ^27^, and this variability has further limited structural characterization. We observe that capsid protein stoichiometry can be modulated by expression-system conditions and harvest time ^28–30^, producing structurally distinct capsid populations that differ in the localization and ordering of VP1/VP2 N-terminal residues. We employed cryo-EM single-particle and subparticle reconstructions to compare empty baculovirus-expressed AAV2 capsids at time points with maximized VP1/VP2 ratios relative to VP3 and time points with VP stoichiometries closer to the conventional 1:1:10 VP1:VP2:VP3 ratio. With these controlled expression conditions, alternative conformations of VP1/VP2 N-termini near the twofold and fivefold axes were resolved, and the relationship between VP stoichiometry and alternative N-terminal configurations was established.

## Results

### VP stoichiometry shifts with harvest time

Based on variations of VP1/VP2/VP3 ratios in prior AAV preparations reported in the literature and observations from our own preparations, ^27,31^ we hypothesized that the relative abundance of the three capsid proteins would change over time during *in vitro* expression. We performed a time-course expression experiment using the baculovirus expression system and collected infected Sf9 cells daily from day 1 through day 5 post-infection. Western blot analysis of aliquots from the time-course experiment (Fig. 1A) was conducted using a monoclonal antibody directed against the C-terminus of VP1, VP2, and VP3. We observed distinct changes in VP expression and relative VP composition throughout the expression time course. On day 1, we detected minimal expression, with only a faint band near 62 kDa, corresponding to VP3. By day 2, band intensity for the three VPs increased significantly. On day 3, VP1, VP2, and VP3 were present at approximately a 2:3:5 ratio, representing the highest overall VP expression and the highest relative abundance of VP1 and VP2, accounting for 17% and 32% of the total VP signal, respectively. By day 4, the relative abundance of VP1 and VP2 declined, and by day 5 the VP1:VP2:VP3 ratio shifted to approximately 1:2:7.

**Fig. 1.**
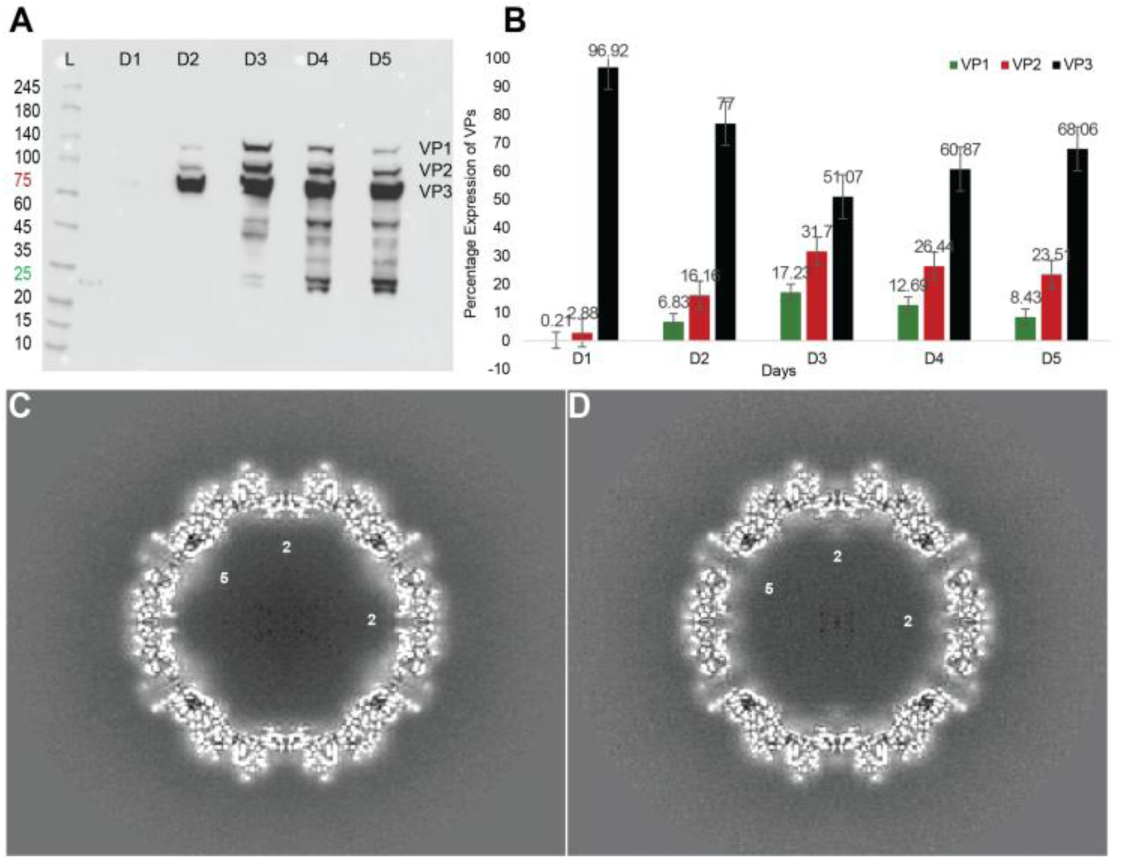
Relative expression of AAV2 VP1, VP2, and VP3 over five days. (A) Western blot analysis of Sf9 cell lysates collected daily from day 1 to day 5 post-infection, probed with a monoclonal antibody targeting the common C-terminus of VP1, VP2, and VP3. Molecular weight markers are shown on the left. (B) Quantification of band intensities expressed as a percentage of the total VP signal per day. The VP1:VP2 ratio is approximately 2:3:5 on day 3 and 1:2:7 on day 5. Bars represent the mean of eight biologically independent experiments (n = 8), and error bars indicate the standard error of the mean (SEM).day (C) Central section through the day 5 cryo-EM map showing diffuse density at the fivefold axis interior and the absence of prominent protrusion density beneath the twofold axis. Symmetry axes are labeled (2 and 5). (D) Central section through the day 3 map showing prominent internal protrusion density beneath the twofold and axis and the absence of organized internal density at the fivefold axis.

Band intensities were quantified using ImageJ across eight biologically independent experiments (n = 8; error bars represent SEM), revealing a statistically significant change in the relative expression of VP1, VP2, and VP3 over the five-day expression time course (Fig. 1B). Post hoc analysis further showed that VP1 and VP2 expressions were significantly higher on day 3 than on all other harvest days. Consequently, we selected day 3 and day 5 for cryo-EM analysis as preparations with distinct VP compositions, representing a VP1/VP2-enriched population and a comparatively VP3-dominant population, respectively.

### Cryo-EM single-particle reconstructions reveal structural differences

Empty AAV2 capsids expressed in Sf9 cells were harvested on day 3 and day 5, purified, and then cryogenically prepared for cryo-EM data collection. The data were subsequently reconstructed using cryo-EM single-particle analysis to resolutions of 2.3 Å and 2.7 Å, respectively, as assessed by Fourier shell correlation at the 0.143 criterion (Fig. S2A). Both maps (Fig. 1C and D) show AAV2’s external icosahedral architecture, including depressions at the twofold axes, protrusions around the threefold axes, and cylindrical pores at the fivefold axes. However, central sections through the two maps reveal notable differences in the distribution of internal densities. In the day 5 map (Fig. 1C), diffuse density is observed within the fivefold pore channel and in the capsid interior near the fivefold axes (labeled 5), with little or no density detected at the twofold axes. In contrast, the day 3 map (Fig. 1D) shows a distinct internal density protrusion at the twofold axes (labeled 2), while the density within the fivefold interior is substantially reduced relative to the day 5 map.

### βA strand organization differs in day 3 and day 5 maps

Examination of the icosahedrally averaged maps revealed substantial differences in βA strand density between the two preparations (Fig. 2). The day 5 reconstruction (EMD-78746) (Fig. 2A) shows continuous, well-ordered density along the βA strand, extending from the βA-βB turn preceding residue 237 to residue 221. By contrast, the day 3 reconstruction (EMD-78754) (Fig. 2B) shows no corresponding density for the βA strand; recognizable structure begins only after residue 237 within the βA-βB turn. This variation in the presence of well-ordered βA density provides a possible structural explanation for the absence of the N-terminal strand in cryo-EM reconstructions of AAV2-AAVR complexes, which has previously been attributed to conformational flexibility in the VP1/VP2 N-terminal regions ^27^.

**Fig. 2.**
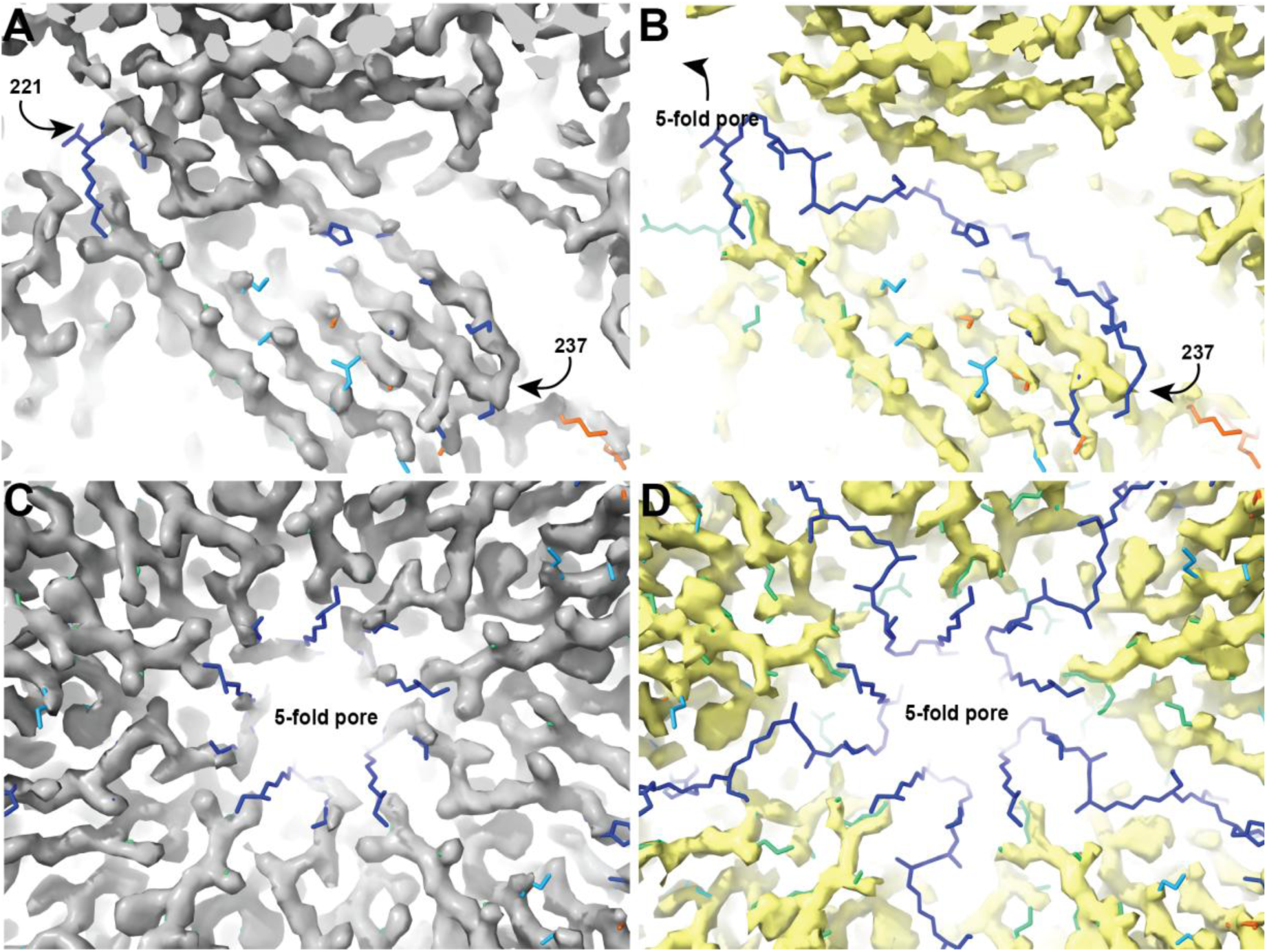
The βA strand is well-resolved at day 5 but disordered at day 3, and the fivefold pore organization differs between preparations. (A, B) The inside of the icosahedral maps near the βA strand is overlaid with a fitted day 5 model (PDB 38DR). The day 5 map (A) is EMD-78746, and the day 3 map (B) is EMD-78754. Panel A shows continuous density for residues preceding residue 237 at day 5 that is absent at day 3. The fivefold pore is labeled in (B). (C, D) Fivefold pore region viewed from the capsid interior for day 5 (C) and day 3 (D); the pore channel is labeled. The day 5 pore is constricted and uniformly organized, whereas the day 3 pore is more open and less densely packed.

The organization of the fivefold pore also varies considerably between the two preparations. In the day 5 map (EMD-78746) (Fig. 2C), the β-ribbon loops surrounding the fivefold channel are uniformly organized, resulting in a constricted pore consistent with other reported AAV2 structures. Conversely, the day 3 map (EMD-78754) (Fig. 2D) shows reduced, less organized density in this region, resulting in a dramatic increase in the fivefold pore opening.

### Asymmetric subparticle reconstruction at the twofold axis resolves an internal protrusion density in day 3

To characterize the internal density observed along the twofold axis in the day 3 icosahedral map, we performed local subparticle reconstruction at the twofold axis using signal subtraction in RELION, followed by refinement without symmetry and by 3D classification (Fig. S4). Central sections through the resulting subparticle reconstructions reveal a striking conformational difference between the day 3 and 5 preparations. The day 5 twofold subparticle map (Fig. 3A) displays no density at the twofold axis within the capsid interior but shows densities around the fivefold corresponding to the internal features observed in the day 5 icosahedral map (Fig. 1C). By contrast, the day 3 twofold subparticle map (Fig. 3B) reveals a compact, well-defined density feature within the capsid interior at the twofold axis. This feature appears as a strong, localized signal distinct from the capsid shell and is significantly more prominent than the corresponding region on day 5 (Fig. 3A,B). This internal density corresponds to the twofold protrusion observed in the day 3 icosahedrally averaged map (Fig. 1D). It aligns with the alternate VP1/VP2 N-terminal configuration previously described in cryo-electron tomography of AAV2-AAVR complexes ^27^. Three-dimensional surface rendering of the twofold subparticle maps confirms these observations. The day 5 map (EMD-79068) (Fig. 3C) shows a clean, unoccupied twofold axis on the inner capsid surface with no protruding feature, whereas the day 3 map with its maximized VP1 and VP2 (EMD-79067) (Fig. 3D) reveals a prominent, globular protrusion extending from the inner capsid surface at the twofold axis. This protrusion connects to the two opposing βA/βB turns and is consistent with alternate N-terminal conformations converging at the start of βB. In this scenario, the protrusion corresponds to residues in and N-terminal to the first strand of the β-sheet (βA), which has been observed at various levels of occupancy and order in prior structures.

**Fig. 3.**
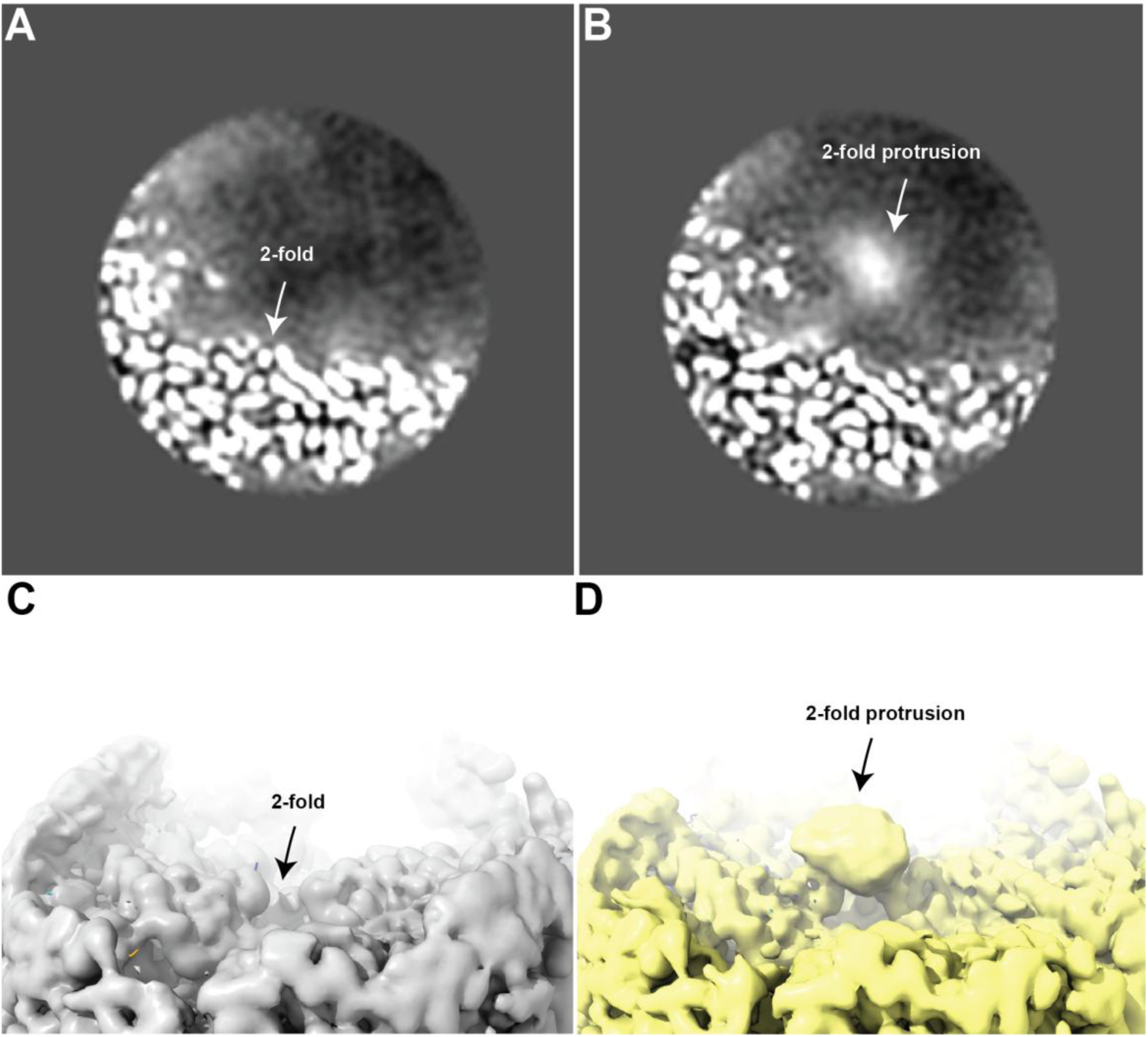
Asymmetric subparticle reconstruction at the twofold axis resolves an internal protrusion density in day 3. (A, B) Central sections through the day 5 (A) and day 3 (B) twofold (local) subparticle reconstructions. The section cuts through the capsid map near the twofold symmetry axis focusing on the interior of the capsid. In day 5 (A), there is no density on the interior surface at the twofold, whereas in day 3 (B), a compact, well-defined feature is observed at the twofold-axis interior (arrow). (C, D) Three-dimensional renderings of the day 5 (EMD-79068) (C) and day 3 (EMD-79067) (D) twofold subparticle maps viewed from the capsid interior. A prominent globular density near the twofold inner capsid surface appears in day 3 (arrow, D) but is absent at day 5 (C). Maps are displayed at equivalent contour levels.

### Subparticle reconstruction at the fivefold axis reveals distinct βA strand configurations

To characterize the alternate conformations of the βA strand, the reconstructed fivefold subparticles were subjected to C5 refinement followed by C5 symmetry expansion (Fig. S5). The expanded fivefold particles were used to generate an asymmetric subunit defined by a 30 Å focused mask generated from a fitted asymmetric subunit model PDB-6IHB ^32^ and applied to both day 3 and day 5 datasets. Classification of the asymmetric subunits for both days was performed without symmetry to resolve differential density within the classes. The day 5 asymmetric subparticle reconstruction (EMD-79069; Fig. 4A) reveals continuous, well-ordered βA strand density extending to residue 219, two residues beyond what could be resolved in the day 5 icosahedral reconstruction (Fig. 2A). By contrast, the day 3 class (EMD-79068; Fig. 4B) shows partial βA strand density traceable only from residue 226, with less organized density at the βA-βB turn, while an additional day 3 class shows a conformation in which βA strand density is not interpretable before residue 237 (Fig. 4C). Together, the three panels demonstrate three discrete βA strand configurations: βA is continuous under VP3-dominant conditions, whereas the VP1/VP2-abundant day 3 preparation contains classes with partial and disordered βA density.

**Fig. 4.**
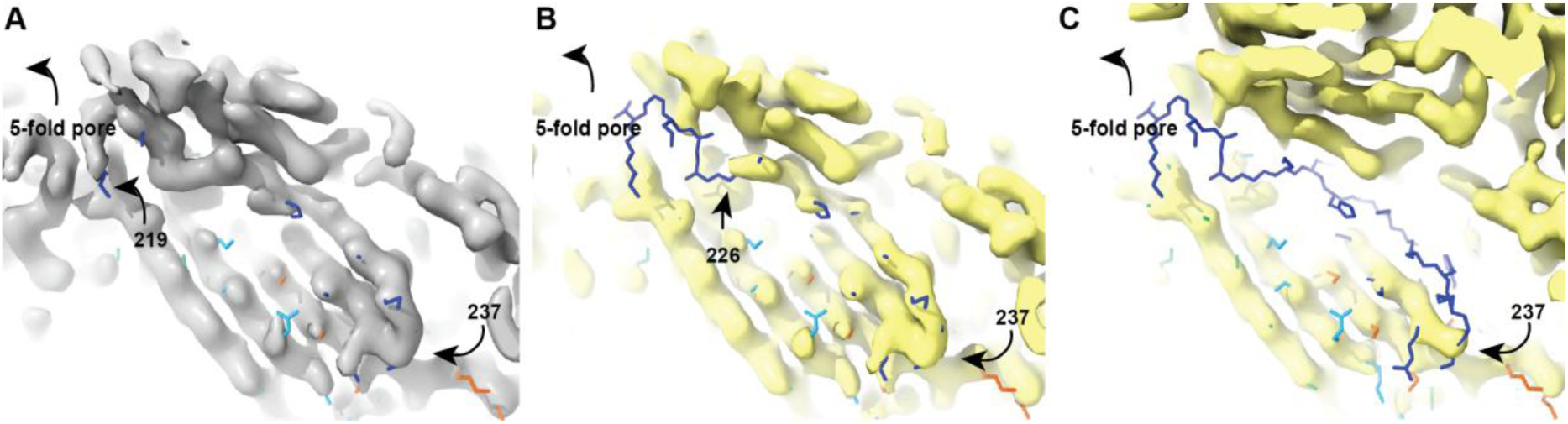
Subparticle reconstruction at the fivefold axis reveals distinct βA strand configurations. (A) The day 5 asymmetric subunit subparticle class showing continuous, well-ordered βA strand density traceable from residue 219 (EMD-79069). (B) A day 3 subparticle class showing partial βA strand density traceable from residue 226 through the βA-βB turn (EMD-79068). (C) A different day 3 class in the same orientation and at the same contour level; lacking interpretable density for βA before residue 237.

### Subparticle reconstruction at the fivefold reveals a plug and internal density around the axis

Beyond the βA strand, the day 5 fivefold subparticle reconstruction overlaid with the day 5 asymmetric subunit (Fig. 5A) reveals a clearly defined density plug occluding the fivefold pore. This feature is absent in the day 3 reconstruction (Fig. 5B). In addition to the plug, internal density is observed surrounding the fivefold axis in the day 5 sample (Fig. 5C). Neither of these features is apparent in day 3 fivefold (Fig. 5D), in which the fivefold pore appears open and unoccupied.

**Fig. 5.**
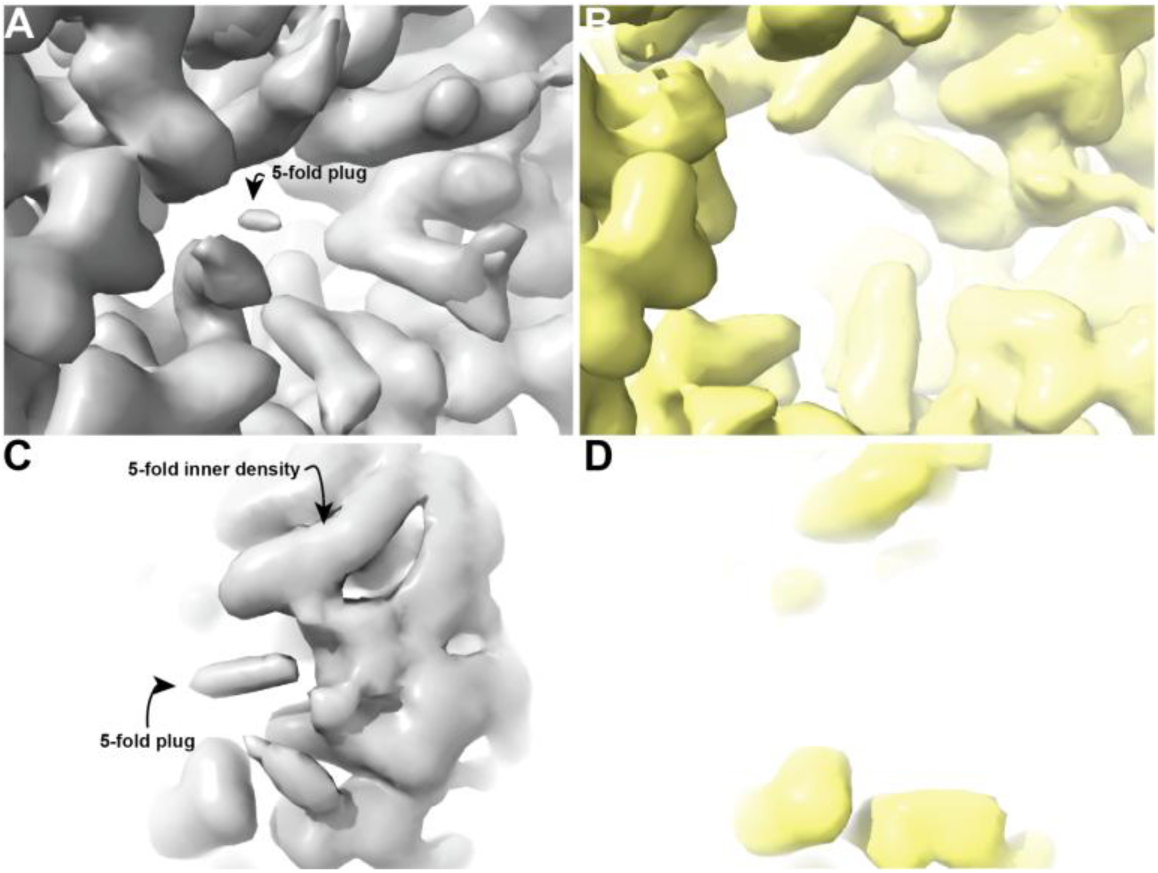
Subparticle reconstruction at the fivefold reveals a plug and internal density around the axis. Day 5 (A) and day 3 (B) subparticle reconstructions of the fivefold pore. Each panel shows an overlay of the complete fivefold pore reconstruction and an asymmetric subunit-focused reconstruction to illustrate the asymmetric unit in the context of the pore. Reconstructions are viewed from inside the capsid along the fivefold axis. A density plug is present along the fivefold axis in day 5 (A) but absent in day 3 (B). The corresponding day 5 (C) and day 3 (D) fivefold asymmetric-subunit maps are shown separately, viewed from the inside of the capsid with the fivefold axis rotated into the plane and at a contour level highlighting the fivefold plug and internal density. These features are apparent in day 5 (C) but are not resolved in day 3 (D).

### Subparticle reconstruction reveals differential ordering of the C-terminal loop

The twofold subparticle reconstructions also revealed differences in the ordering of a C-terminal loop spanning residues 703-710. In day 5 icosahedral reconstruction (Fig. S3A), this loop is clearly defined and continuous, whereas in the day 3 icosahedral map (Fig. S3B), it appears disordered. Examination of the corresponding twofold subparticle reconstructions for day 5 (Fig. S3C) and day 3 (Fig. S3D) further resolved the loop density. In day 3 reconstruction, density for residues 703-710 is resolved, revealing a distinct loop conformation that is not apparent in the icosahedral map. Although density for this loop is already present in day 5 icosahedral map, the twofold subparticle reconstruction further resolves a more defined loop conformation (Fig. S3C).

## Discussion

Despite decades of studies, the structure of the N-termini of AAV2 capsid proteins VP1, VP2, and VP3 has remained elusive largely due to conformational heterogeneity, symmetry mismatch, and the low relative stoichiometry of VP1 and VP2 relative to VP3, which, together, have limited the resolution of the N-terminal region ^10,27,33,34^. Consequently, high-resolution structures of AAV2 determined to date have resolved only the β-jellyroll fold that is common to all three proteins. The unique N-terminal residues of VP1 (VP1u), the common N-terminal residues of VP1 and VP2 (VP1/VP2 common region), and approximately the first 20 amino acids of VP3 have remained structurally undefined ^33^. Furthermore, unaccounted features in prior cryo-EM maps, attributed to the N-terminal region, lay on a two-fold symmetry axis, obfuscating detailed interpretation ^10,33,34^. By analyzing AAV2 viral-like particles at time points with distinct VP1/VP2/VP3 compositions and applying cryo-EM subparticle reconstruction, we resolved heterogeneous N-terminal conformations that were obscured in prior icosahedrally averaged single-particle reconstructions.

In the day 3 preparation where VP1 and VP2 accounted for 50% of the total VP content, subparticle reconstructions revealed partially ordered density protruding from the inner capsid surface near the twofold symmetry axis (Fig. 3). This density likely arises from the VP1/VP2 N-terminal regions and is consistent with previous reports attributing diffuse density near the twofold axes to these regions^11,20,27,35^. Our observations suggest that differences in VP stoichiometry may explain some of the discrepancies in βA strand organization reported in previous studies^28,34,36,37^. Focused classification of our day 3 and day 5 maps revealed that VP1/VP2 enriched capsids contain multiple conformations, with some subunits adopting the canonical βA conformation and others exhibiting an alternative VP1/VP2 N-terminal conformation in which density extends inward along the twofold axis from the βA-βB turn of the two opposing asymmetric units.

Classification of the day 3 fivefold subparticles revealed two structural classes distinct from the day 5 reconstruction. Whereas the day 5 reconstruction exhibited continuous βA density extending from residue 219 and connecting to a ring of density partially occluding the fivefold pore, the day 3 classes showed less ordered βA density extending from residue 226 or no interpretable density before residue 237. Neither day 3 class exhibited the fivefold-proximal density observed in day 5. These differences suggest that increased VP1/VP2 abundance promotes conformational heterogeneity of the N-terminal regions, potentially by disrupting VP3 interactions near the fivefold axis. We propose that this disruption favors alternative conformations in which a greater fraction of N-terminal extensions are directed toward the capsid interior rather than contributing to the ordered βA strand and fivefold-proximal density.

Our observations suggest a model for VP1/VP2 externalization that is consistent with the structural heterogeneity observed in day 3 and day 5 preparations. In wild-type (infectious) AAV2, the approximately 1:1:10 stoichiometry of VP1:VP2:VP3 means that the majority of βA strands and the ring of density surrounding the fivefold pore must be contributed mostly by VP3 ^21,33,38^. In our model, we would predict that βA for the remaining capsid proteins would be disordered. During endosomal trafficking, AAV2 capsids undergo a critical conformational change thought to be triggered by the acidic endosomal environment, exposing the N-terminal portions of VP1 and VP2, which are initially sequestered within the capsid interior ^39,40^. This mechanism is consistent with cryo-EM structures of the AAV9 capsid, which show that progressively acidic pH (7.4 to 4.0) increasingly occludes the fivefold channel with a density column that likely corresponds to part of a glycine-rich segment starting within the VP1-2 common region and ending after the start of VP3 ^41^. Higher fractions of the bulkier VP1/2 N-terminal regions in day 3 may be harder to accommodate within the fivefold structures, which could lead to their destabilization. Increasing VP1 and VP2 could increase the incidence of alternative conformations of the βA strand. One could also speculate that increasing fractions of VP1 or VP2 extruded at low pH might facilitate accommodation of capsid proteins that remain internal in configurations compatible with the ordered βA. Thus, the structural differences documented in the current work may be linked to AAV’s conformational transition required for endosomal escape and nuclear localization. Future research is needed to characterize the exact mechanism by which these regions are exposed for endosomal escape and nuclear localization.

## Materials and Methods

### Cell Culture

Empty AAV2 virus-like particles were produced in Sf9 cells using the Invitrogen Bac-to-Bac baculovirus expression system, following established protocols ^42–45^. Sf9 insect cells, derived from *Spodoptera frugiperda* ovaries, served as the host for baculovirus-mediated capsid protein production ^46^. Master and working cell stocks were cultured under sterile conditions following the Bac-to-Bac protocol (MAN0000414), grown in suspension in protein-free ESF 921 Insect Cell Culture Medium (Expression Systems, Davis, CA, USA; Cat. NC9541308) at 27°C with gentle shaking (100-130 rpm). Cell density and viability were monitored daily using trypan blue exclusion and an automated Countess cell counter (Invitrogen), with passaging to maintain logarithmic growth at 1-2 × 10⁶ cells/mL and viability above 95%. Confirmation of cell density and viability was essential before proceeding to transfection.

### AAV2 Capsid Protein Expression

To produce AAV2 capsid proteins, recombinant bacmid DNA ^42^ was used to transfect log-phase Sf9 cells using Cellfectin II reagent according to the Bac-to-Bac protocol. The bacmid DNA (1 μg/mL) was mixed with Cellfectin (Gibco/Thermo Fisher Scientific, Grand Island, NY, USA; Cat. 10362100) in Grace’s Insect Media-Supplemented (Gibco/Thermo Fisher Scientific, Grand Island, NY, USA; Cat. 11605094), allowed to form complexes, and then added to cells in 6-well plates. After 5-6 hours, the medium was replaced with fresh Sf-900 II SFM, and cells were incubated at 27°C for 3-5 days for viral replication and protein expression. The P1 viral stock was harvested from the supernatant and clarified by low-speed centrifugation (100 x g). For amplification, fresh Sf9 cells were infected with P1 at an MOI of 0.1-1 and incubated for 72 hours, yielding the high-titer P2 stock, which was stored at 4°C for future use.

### Time Course Experiment

Time-course expression studies were conducted over five days post-P1 infection using ten 125 mL Erlenmeyer flasks with 50 mL of baculovirus-infected Sf9 cell culture. We monitored VP protein levels, cell viability, and total count daily. Each day, 10 mL of culture was collected, centrifuged at 1,000 × g for 10 minutes, and the pellet was resuspended in 1 mL of 1× PBS for storage at -80°C until Western blot analysis. For lysis, 200 μL of the resuspended pellet was diluted in 800 μL of lysis buffer, including 150 mM NaCl, 20 mM Tris-HCl, 0.5% CHAPS, 0.5% sodium deoxycholate, with protease inhibitors (Pierce Biotechnology/Thermo Fisher Scientific, Rockford, IL, USA; Cat. PIA32963) and Benzonase (Merck KGaA, Darmstadt, Germany; Cat. 1016540001). Samples were lysed at room temperature with gentle rocking for 1 hour. A 40 μL aliquot of the lysate was mixed with 40 μL of 2× Laemmli loading dye, heated at >95°C for 10 minutes, and loaded onto a 4-20% SDS-PAGE gel (GenScript, Piscataway, NJ, USA; Cat. M42015) in Tris-MOPS-SDS Running Buffer (GenScript, Piscataway, NJ, USA; Cat. M00138). Proteins were separated at 140 V for 1 hour and 30 minutes and transferred to a nitrocellulose membrane at 45 V for 45 minutes. The membrane was blocked in 5% BSA in TBS overnight and incubated for 1 hour with MonoRab™ AAVX VP1/VP2/VP3 antibody (clone 5G4; GenScript, Piscataway, NJ, USA; Cat. No. A02206). After four washes in Tris-buffered saline with Tween 20 (TBST), it was incubated with anti-rabbit secondary antibody (Rockland Immunochemicals, Limerick, PA, USA; Cat. No. 18-8816-33), washed again, and developed using Immobilon Chemiluminescent HRP Substrate. Imaging was performed with a ChemiDoc Imaging System. Band intensities for VP1, VP2, and VP3 were quantified using ImageJ (National Institutes of Health, Bethesda, MD, USA) from eight independent biological experiments. For each harvest day, the intensity of each VP band was normalized to the total VP signal (VP1 + VP2 + VP3) and expressed as a percentage of the total capsid protein signal. This percentage reflects relative band intensity. Data are presented as the mean ± standard error of the mean (SEM). Statistical analyses were performed separately for VP1, VP2, and VP3 using one-way analysis of variance (ANOVA) in Microsoft Excel. When a significant main effect was detected, Tukey’s honestly significant difference (HSD) post hoc test was performed in JASP (Version 0.97) to identify pairwise differences among harvest days. Statistical significance was defined as *p* < 0.05.

### AAV2 Capsid Protein Production: Lysis

For large-scale capsid protein production, Sf9 cells from 500-1000 mL cultures harvested on day 3 and day 5 were pelleted and stored at -80°C prior to purification. On purification day, cell pellets were thawed, resuspended in 100 mL of sterile lysis buffer (150 mM NaCl, 20 mM Tris-HCl, pH 8.0), and lysed by adding 12.5 mL of 10% (w/v) sodium deoxycholate (MilliporeSigma [Sigma-Aldrich], St. Louis, MO, USA; Cat. D6750). To reduce viscosity, 27 μL of Benzonase nuclease (Merck KGaA, Darmstadt, Germany; Cat. 1016540001) was added, and the mixture was vortexed every 10 minutes for 1 hour at 37 °C. Cellular debris was removed by centrifugation at 30,000 × g for 60 minutes at 4°C, and the supernatant was filtered through a 0.45 μm PVDF syringe filter ^47^. An aliquot of the clarified lysate was retained for quality analysis.

### Purification of Lysate

The clarified lysate was purified using heparin affinity chromatography on an FPLC system at 4°C. The system was washed and equilibrated according to standard protocols using 20% ethanol and ultrapure water. A prepacked 5 mL heparin column was equilibrated with ultrapure water, buffer A (100 mM NaCl, 20 mM Tris-HCl, pH 8.0), and buffer B (500 mM NaCl, 20 mM Tris-HCl, pH 8.0). The lysate was loaded at 0.5 mL/min after pre-equilibration with 12.5% buffer B. The column was washed with 20 column volumes of buffer A, then eluted using a gradient to 50% buffer B over 5 column volumes, followed by step elution to 90% and 100% buffer B, each for 5 column volumes. Fractions were collected in 1 mL aliquots and stored at -20°C ^47^.

### Concentration of Viral Particles

Pooled FPLC fractions were concentrated using 100 kDa molecular weight cutoff centrifugal filter units (Amicon Ultra, MilliporeSigma). The pooled fractions in a 50 mL unit were centrifuged at 2,000 × g for 5 minutes, reducing the volume to approximately 500 μL. The concentrated sample was then transferred to a 15 mL unit (100 kDa molecular weight cutoff, MilliporeSigma) for buffer exchange, where 1 mL of sterile PBS was added and gently mixed. Centrifugation at 2,000 × g for 2 minutes yielded approximately 100 μL of concentrated, purified AAV2 virus-like particles suitable for cryo-EM grid preparation, as determined by NanoDrop absorption.

### Cryo-EM Sample Preparation and Data Collection

Quantifoil R1.2/1.3 300-mesh gold grids (Quantifoil Micro Tools GmbH, Jena, Germany; Cat. Q425CR1.3) were glow-discharged for 20 seconds using a Solarus 950 plasma cleaner before sample application. For the day 3 sample, 4 μL of AAV2 VLP at 1.34 mg/mL was applied to the glow-discharged grid and vitrified using a Vitrobot Mark IV at 4°C and 100% humidity, with a blot force and time of 1 and 1 second, respectively, by plunge-freezing in liquid ethane cooled by liquid nitrogen. The day 5 sample was prepared similarly at a concentration of 1.22 mg/mL. The vitrified grids were clipped into autogrids using a cryo-clipping station and stored in liquid nitrogen until data collection

Data collection was conducted on a Titan Krios transmission electron microscope at 300 kV, using a DE Apollo direct electron detector. Movies were captured with Leginon ^48^ at 47,000× magnification and a pixel size of 0.962 Å, with defocus ranges of -1.00 to -3.00 μm for both days. The total electron dose was 59.59 e⁻/Å² over 113 frames for day 3 and 59.98 e⁻/Å² over 114 frames for day 5.

### Image Preprocessing and Reconstruction

All movie frames were aligned and dose-weighted using MotionCor2 ^49^. Contrast transfer function (CTF) parameters for the motion-corrected micrographs were estimated with CTFFIND-4.1 ^50^. For day 3, 121 particles were manually picked from 3,259 micrographs to train the Topaz particle picker ^52^ at a 260 Å particle diameter, resulting in 51,475 particles. After 2D classification, 32,304 particles were advanced to 3D classification without further alignment and using a reference AAV2 map in RELION 5 ^53,54^. This process retained 21,228 particles for 3D auto-refinement, enforcing icosahedral symmetry. A soft mask was generated using a binarization threshold of 0.003, a low-pass filter of 15 Å, and a 3-pixel soft edge and binary extension, yielding a final reconstruction at 2.3 Å resolution. Similarly, day 5 recovered 178 particles from 2,416 micrographs to train Topaz, yielding 784,686 particles extracted using a 448-pixel box size and rescaled to 224 pixels in RELION 5. After 2D and 3D classification, 17,900 particles were selected for 3D auto-refinement. CTF refinement, beam-tilt correction, and a soft mask (binarization threshold of 0.005) yielded a final reconstruction at 2.7 Å resolution.

## Subparticle Reconstruction

### Geometric Reconstruction of the Twofold and Fivefold

A spherical mask with a 100 Å radius was created in ChimeraX using the "shape sphere" command, centered on a single fivefold pore. The vector from the capsid’s center of mass to the sphere center was calculated, and the remaining 11 fivefold directions were determined by applying icosahedral symmetry operators to this unit vector, resulting in 12 sphere centers for all fivefold axes. Sphere placement was visually inspected and confirmed in ChimeraX (Fig. S5).

Twofold axis masks were generated using the same geometric procedure as the fivefold axes. Using the day 5 map as a reference, a spherical mask was defined in ChimeraX, centered on one twofold axis with a radius of 100 Å, similar to the fivefold reference sphere. The center of mass of the capsid map and the center of the twofold reference sphere were measured in scene coordinates to compute the twofold direction vector and unit vector, resulting in 30 twofold sphere centers (Fig. S4). All twofold spheres were converted into volume masks in ChimeraX and exported as MRC files into RELION.

### Masks Creation and Signal Subtraction

We converted the fivefold and twofold spherical MRC masks generated in ChimeraX into soft masks with a 3-voxel extension and a 3-voxel soft edge in RELION. Following mask import, signal subtraction was performed referencing the *optimiser.star* and *data.star* files from the 3D auto-refined jobs. The resulting subparticles were written with a 256-box size. The process was repeated for each symmetry-related mask and applied to both day 3 and day 5 datasets.

An initial reference for refining the signal subtracted particles was made by applying one of the soft spherical masks from the previous step to the unsharpened full-capsid map using *relion_image_handler* to create a masked volume. The center of mass of the masked region was calculated in RELION, yielding a shift vector (Δx, Δy, Δz) indicating its displacement from the particle box center. This vector was then applied with the opposite sign using *relion_image_handler* to recenter the local density at the signal-subtracted box origin and cropped to 256³ voxels.

### Subparticle Processing at the Fivefold Axis

The refinement of the day 3 and day 5 fivefold signal-subtraction was conducted using C1 symmetry (Fig. S5). The C1 map had a plug in the middle of the fivefold pore. To better characterize this density, we performed a C5 refinement, followed by symmetry expansion and classification. A focused mask around the asymmetric subunit was created by fitting one subunit into the day 5 C5-refined map and converting the subunit into a binary mask in ChimeraX. The mask was then processed in RELION with low-pass filtering, binarization, a 3-pixel binary extension, and a 3-pixel soft edge, resulting in a soft mask of a single subunit applicable to both time points. Focused 3D classification was performed on the C5 symmetry-expanded particles under C1 symmetry to classify each symmetry-related particle based on the density within the asymmetric unit.

### Subparticle Processing at the Twofold Axis

For the twofold analysis, the clipped map with a 256-pixel box served as the reference map (Fig. S4). Subtracted particles from days 3 and 5 were generated separately and subjected to 3D refinement without symmetry using the same settings as the fivefold refinement. The resulting Refine3D volume was imported into ChimeraX to create a focused mask with a 50 Å radius centered on the twofold axis. This spherical mask was converted to a volume mask with the ChimeraX, exported as an MRC file, and softened in RELION using the same binarization, extension, and soft-edge parameters as the fivefold mask. The resulting soft twofold mask was used as the solvent mask for 3D classification without alignment, with the Refine3D volume serving as the reference map and the corresponding star file as input.

### Model Generation

Initial coordinates were derived from three reference structures: AAV2 atomic model PDB 6IHB ^32^, AAV-DJ structure PDB 7KFR (1.56 Å) ^55^, and AAV2 structure PDB 6E9D (1.9 Å) ^56^. We performed preliminary modeling and refinement in the presence of solvent water. Two translations were applied to align the coordinates with the map origin and to correct a minor pixel-size discrepancy. Symmetry-related neighboring residues were generated for context during manual rebuilding of residues 219 through 735 in the program Coot ^57^ guided by prior structures PDB 7KFR and PDB 6E9D. Rotamer corrections and backbone adjustments were made where the experimental density indicated deviations from the reference structures. Real-space refinement was performed using an updated implementation of RSRef ^58^ that uses the Amber ^59^ force field in OpenMM ^60^ for stereochemical restraint and treats solvent water molecules as TIP3P ^61^ rigid bodies. B-factor refinement followed coordinate refinement.

## Supplemental Figures

**Fig. S1.**
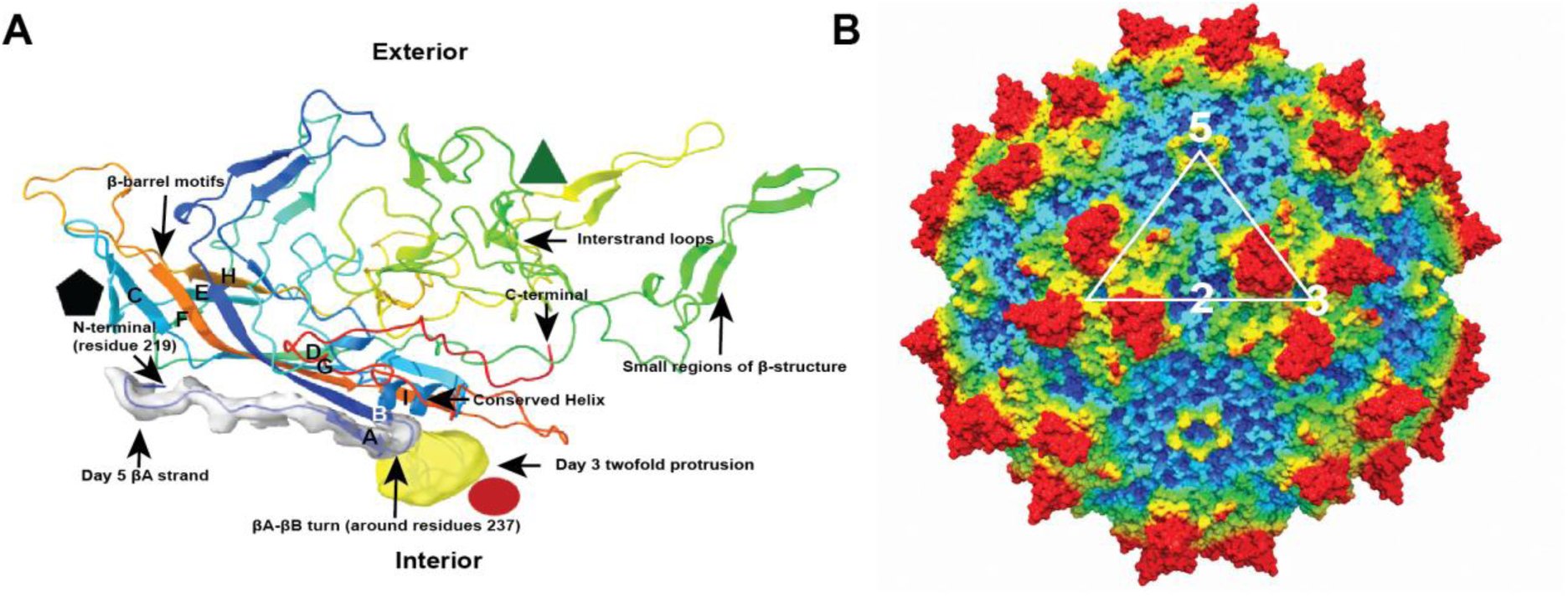
Unique features of the day 3 and day 5 structures in the context of the AAV capsid subunit topology. The ribbon representation of a single AAV2 capsid subunit (PDB 38DR) is colored from N- to C-terminus (blue to red) and shows the nine-stranded β-barrel motif (labeled strands βA-βI). The approximate orientations of the icosahedral twofold (red oval), threefold (green triangle), and fivefold (black pentagon) axes are indicated in A, with strokes showing the approximate directions of the symmetry axes in B.

**Fig. S2.**
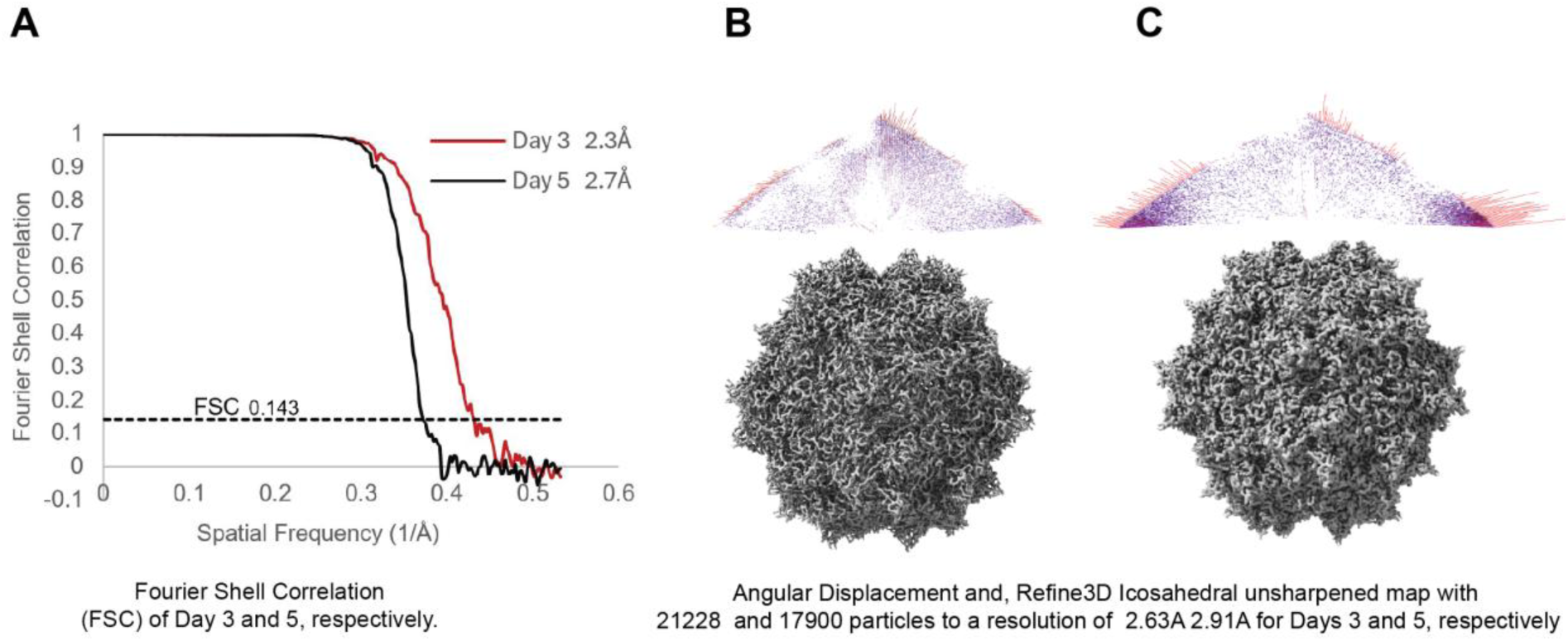
Data quality and resolution assessment for day 3 and day 5 icosahedral reconstructions of empty AAV2 capsids. **(A)** Fourier shell correlation (FSC) curves for day 3 (red, 2.3 Å) and day 5 (black, 2.7 Å) reconstructions, plotted against spatial frequency. The dashed line marks the FSC = 0.143 criterion used to assign the reported resolutions. **(B and C)** Angular distribution of particle orientations assigned during 3D auto-refinement in RELION, shown as a polar plot in which the length and color of each spoke indicate the relative number of particles assigned to that view, and the unsharpened Refine3D icosahedral map generated from the 21,228 particles on day 3 and 17,900 particles on day 5.

**Fig. S3.**
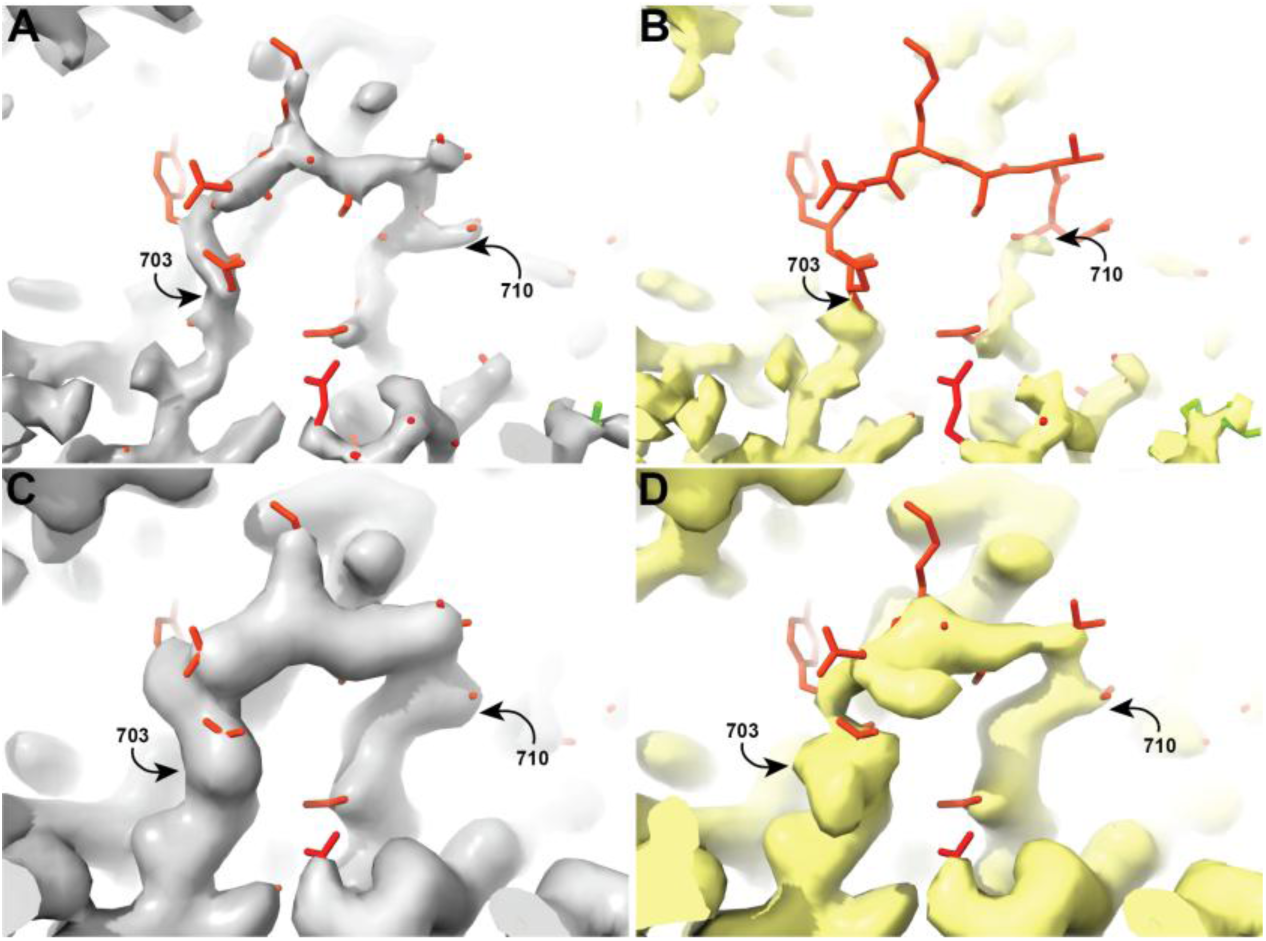
Subparticle reconstruction reveals differential ordering of the C-terminal loop. Map density and fitted model 38DR for residues 703-710 in day 5 global map (A) and day 3 global map (B). The backbone trace is continuous at day 5, but residues 703-710 are missing at day 3. Day 5 (C) and day 3 (D) show subparticle reconstructions within spheres centered on two-fold axes. Focused subparticle averaging recovers density for residues 703-710 in both preparations, with more contiguous density for day 5 than day 3. The maps are displayed at equivalent contour levels.

**Fig. S4.**
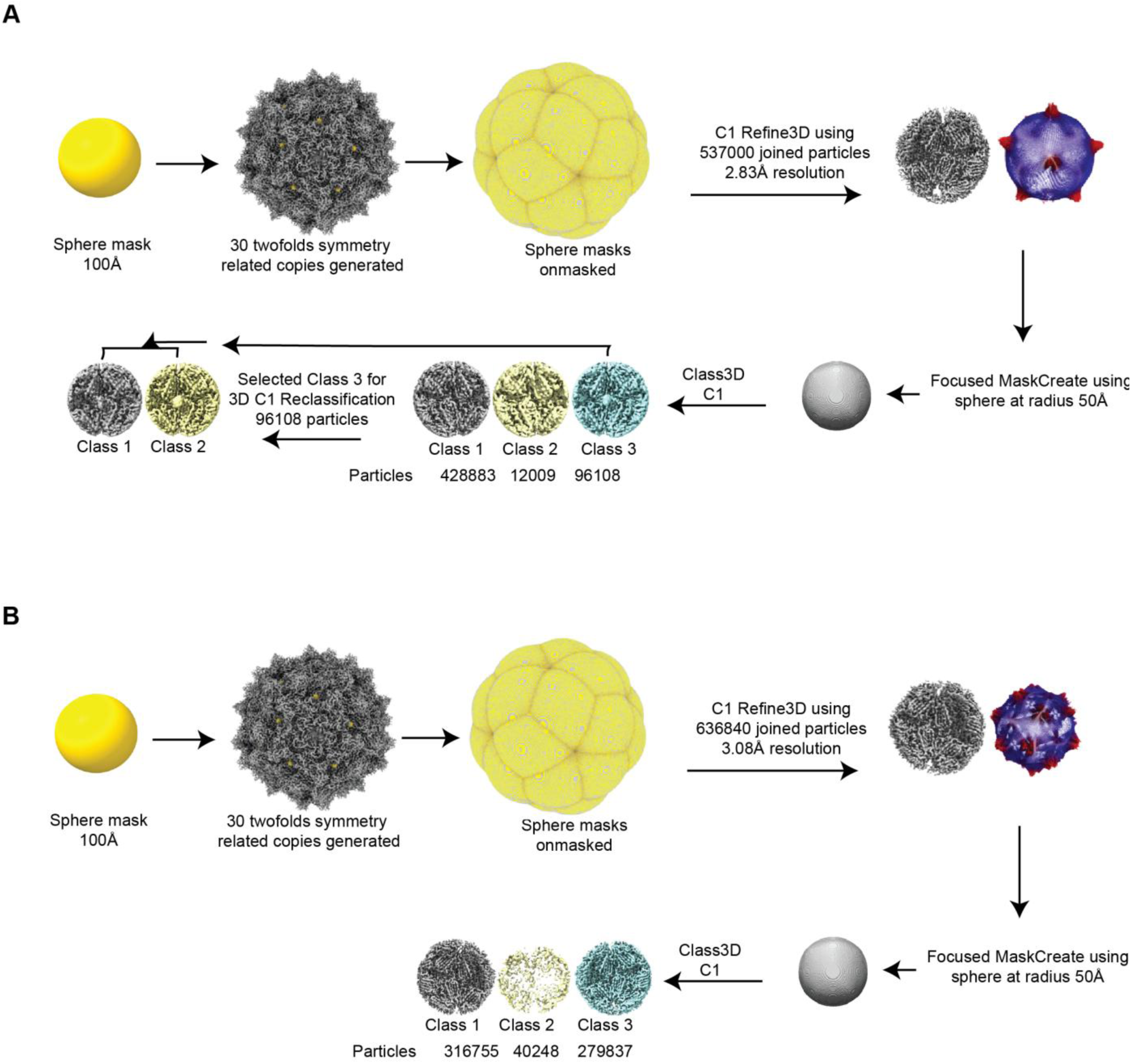
The twofold subparticle reconstruction workflow for day 3 (A) and day 5 (B). Twofold subparticle reconstruction workflow for day 3 (A) and day 5 (B). A spherical mask (100 Å radius) centered on a twofold axis was expanded to 30 symmetry-related copies using icosahedral symmetry operators in ChimeraX and converted to soft masks in RELION. C1 3D auto-refinement of the joined, twofold-subtracted particle sets; 537,000 particles (day 3) and 636,840 particles (day 5), yielded reconstructions at 2.83 Å and 3.08 Å resolution, respectively. A focused mask (50 Å radius) centered on the twofold axis was then used for reference-free focused 3D classification (Class3D, C1 symmetry), sorting particles into three classes (Class 1-3) for each dataset. For the day 3 dataset (A), Class 3, enriched in the twofold protrusion density described in the main text (Fig. 4), was selected for a second round of focused C1 3D classification, yielding 96,108 particles that resolved into two subclasses (Class 1 and Class 2) representing distinct conformational states of the twofold density.

**Fig. S5.**
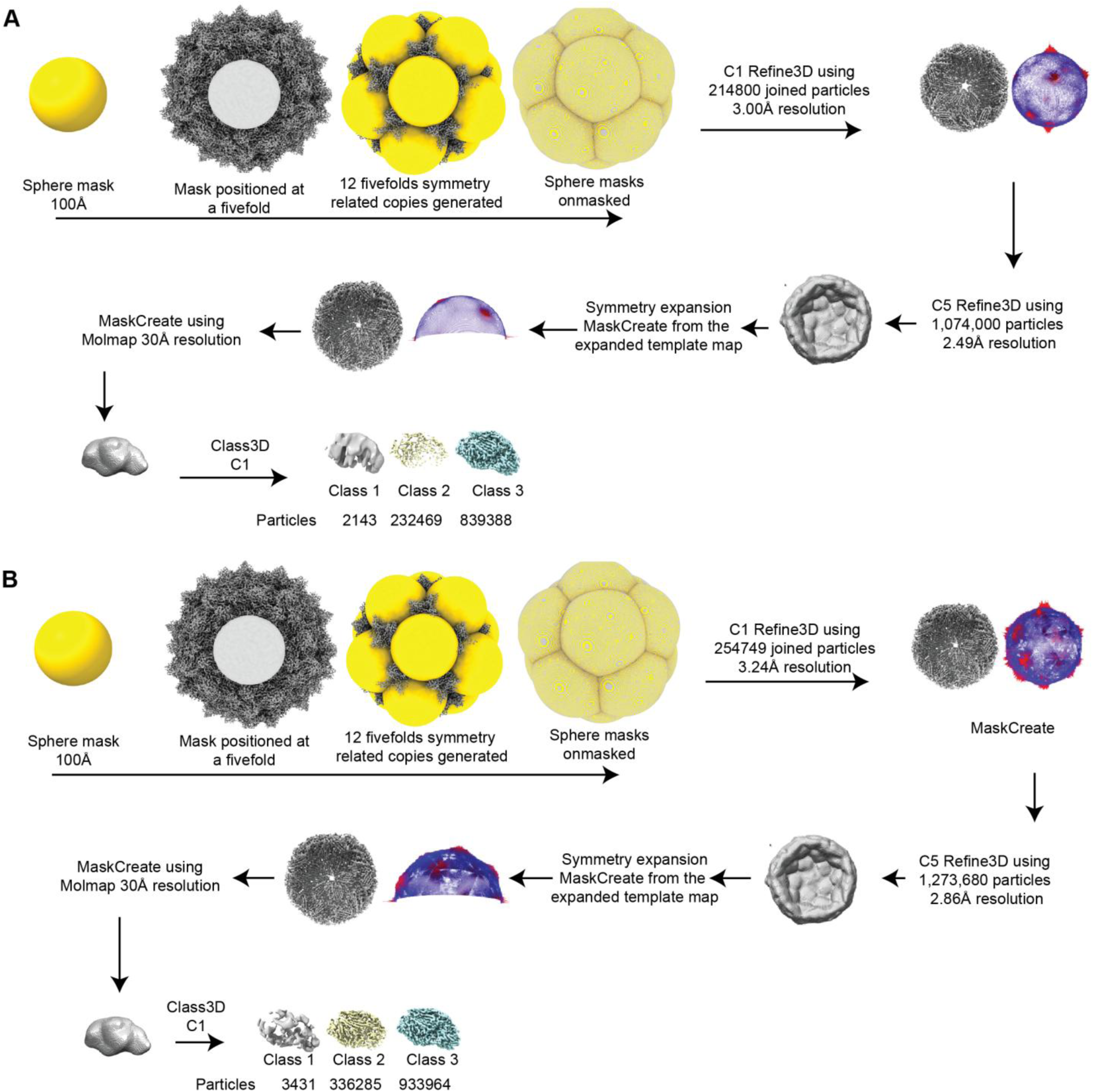
Fivefold subparticle reconstruction workflow, Days 3 (A) and 5 (B) datasets. A spherical mask (100 Å radius) was positioned at a single fivefold axis in ChimeraX, and the remaining 11 symmetry-related copies were generated by applying icosahedral (I1) symmetry operators, yielding 12 fivefold-centered masks that were converted to soft masks in RELION. C1 3D auto-refinement of the joined particle sets; 214,800 particles (day 3) and 254,749 particles (day 5), produced icosahedral reconstructions at 3.00 Å and 3.24 Å resolution, respectively. Symmetry expansion of the refined particles, followed by MaskCreate from the expanded template map, generated a fivefold-subunit reference used for C5 Refine3D of the expanded particle stacks (1,074,000 particles, day 3; 1,273,680 particles, day 5), yielding subparticle reconstructions at 2.49 Å and 2.86 Å resolution. A focused mask around a single fivefold subunit was generated in ChimeraX by molmap simulation at 30 Å resolution and converted to a soft mask in RELION (MaskCreate). Focused 3D classification without alignment (Class3D, C1 symmetry) of the C5 symmetry-expanded particles then sorted individual fivefold subunits into three classes (Class 1-3) based on the density within each subunit.

## Acknowledgements

We thank the National Institutes of Health/National Institute of General Medical Sciences (NIGMS) for funding support R35GM122564 (M.S.C.; S.M.S.). We thank Dr. Gwi Moon Seo and the Institute of Molecular Biophysics Protein Expression Facility (IMB-PEF) at Florida State University for protein expression support. Cryo-EM data were collected at the Biological Science Imaging Resource (BSIR) with support from Dr. Xiaofeng Fu. The Florida State University Research Computing Cluster (HPC) provided computational resources. We thank the Stagg and Chapman laboratories for productive discussions and support throughout this work. Conceptualization and methodology: S.M.S. Funding acquisition: M.S.C and S.M.S. Atomic model building and refinement: M.S.C. and I.O.O. Sample preparation and data collection: A.S., S.K., R.P., TSA, AMA, and I.O.O. Data analysis: I.O.O. and S.M.S. Writing, original draft preparation: I.O.O. and S.M.S. Writing, review and editing: all authors.

